# GRanges: A Rust Library for Genomic Range Data

**DOI:** 10.1101/2024.05.24.595786

**Authors:** Vince Buffalo

## Abstract

**Motivation:** The Rust programming language is a fast, memory-safe language that is increasingly used in computational genomics and bioinformatics software development. However, it can have a steep learning curve, which can make writing specialized, high performance bioinformatics software difficult.

**Results:** GRanges is a Rust library that provides an easy-to-use and expressive way to load genomic range data into memory, compute and process overlapping ranges, and summarize data in a tidy way. The GRanges library outperforms established tools like plyranges and bedtools.

**Availability:** The GRanges library is available at https://github.com/vsbuffalo/granges and https://crates.io/crates/granges.

## Introduction

Genomic data analysis often requires processing *genomic range* data, which is any quantitative data associated with regions or positions relative to a reference genome coordinate system. For example, an RNA-seq analysis may rely on counting the number of mapped reads that overlap a particular exon, and many population genomic analyses require statistical summaries of variation data grouped by genomic feature such as finding the site-frequency spectrum of four-fold degenerate sites. Generic genomic range operation tools like bedtools and bedops [12, 14, 13] implement an essential set of these range operations, and are written in highly performant compiled languages such as C and C++.

However, constructing genomic data analysis pipelines requires that these generic command-line tools are patched together into custom scripts or with workflow managers such as Snakemake [7]. While this approach is flexible and powerful, there is an inherent tradeoff. Since these tools accept plaintext bioinformatics formats (e.g., BED, GFF/GTF, VCF) as input and output, each tool in a processing pipeline must deserialize and serialize the data into and out of memory. This can lead to unnecessary computational overhead and disk space usage. Additionally, these tools lack flexibility; only a subset of statistical functions are supported when summarizing the numeric data associated with overlapping ranges.

By contrast, genomic range processing libraries like Bioconductor’s GenomicRanges [8] are flexible and intended for interactive data analysis. The plyranges package extends GenomicRanges and provides a powerful grammar for manipulating genomic range data inspired by the tidyverse approach to analyzing data [9, 17, 18]. In particular, genomic range operations are expressed in terms of different types of *joins*, since they are fundamentally equivalent to database join operations. While the plyranges interface is a powerful, flexible, and expressive way to develop specialized genomic data analyses, it is implemented in the R language (with C extensions), which imposes some performance tradeoffs compared to compiled software like bedtools and bedops.

Here, I introduce *GRanges*, a Rust-based genomic ranges library and command-line tool for working with genomic range data. The goal of GRanges is to strike a balance between the expressive grammar of plyranges, and the performance of tools written in compiled languages. The GRanges library has a simple yet powerful grammar for manipulating genomic range data that is tailored for the Rust language’s ownership model. Like plyranges and tidyverse, the GRanges library develops its own grammar around an *overlaps-map-combine* pattern. This grammar lowers the barrier to entry for the development of bespoke genomic range processing tools suited to specific analyses and processing tasks, while still producing high-performance compiled software. Even though primarily intended as a library, GRanges also has a command-line tool that implements some of the core functionality of bedtools as an example of how to use the GRanges interface, for benchmarking, and integration tests.

## Rust and the GRanges Generic Types Design

Rust is a compiled, statically-typed, memory-safe programming with speeds comparable to C and C++ [5]. Increasingly, the Rust language is used for bioinformatics and genomics software [10] such as in Rust-Bio [6] and NanoPack2 [3]. Rust ensures memory-safety through its *ownership model*, which enforces that each object in memory has a single owner and memory is freed when the owner goes out of scope. While this design prevents many runtime errors due to bugs in handling memory correctly, it can impose a steep learning curve to new Rust developers.

The interface of the GRanges library is designed to lower such barriers for new developers writing computational genomics software in Rust. The basic type used in GRanges is the `GRanges<R, T>`, generic type, which stores a *range container type* `R` and a *data container type* `T`. The generic range container design allows for different, specialized data structures to be used for different tasks. When loading in genomic ranges, GRanges uses a standard dynamically growing `VecRanges` type, which is the most effective data structure for this task. However, when the overlaps needs to be found between two different `GRanges` objects, different data structures are needed for performance reasons. In this case, the GRanges library uses the Rust `coitrees` library [4] which uses cache-oblivious B-trees [1]. The GRanges library provides the `GRanges::into coitrees()` method to convert `VecRanges` to `COITrees` types.

The `GRanges<R, T>` is also generic over its data container type. This design allows for data to be stored in dynamically growing data containers like Rust’s `Vec<D>` for when it is first loaded in, as well as arrays and matrices in libraries like `ndarray`. In the latter case, storing data in these array-type containers is more amenable to numeric and statistical computing operations, since these operations can be done on array types directly. Each range in the range container contains an index to its corresponding data in the data container. There is no restriction that this relationship must be one-to-one, meaning that memory can be saved by having ranges pointing to the same element if the underlying data is the same. Furthermore, a benefit of the generic statically-typed design of `GRanges<R, T>` is that operations that cannot be computed on particular range or data container types will be known at development or compile time rather than at runtime, which reduces the risk of runtime errors common in dynamically-typed languages like Python.

## The Overlaps-Map-Combine Pattern

Inspired by the plyranges and tidyverse libraries, GRanges uses a method-chaining approach to build up various genomic range and data processing steps. Like plyranges, GRanges’s grammar frames various genomic overlap operations as “joins”, and all downstream processing of overlapping ranges is done by applying or “mapping” a *combine* function to these joins with `GRanges::map joins()`. The combine function takes the joins and their data and combines them into a single summarized data entry. For example, `bedtools map` lacks the ability to calculate a mean of the BED score column weighted by the degree of overlap. This is often disadvantageous for statistical analyses; suppose range A with score 4.1 overlaps a window by 2 basepairs, and range B with score 0.1 overlaps the same window by 50 basepairs. The unweighted mean score for the window would be (4.1 + 0.1)*/*2 = 2.1 but the weighted mean would be 2*/*52 *×* 4.1 + 50*/*52 *×* 0.1 *≈* 0.254. A mean score weighted by the degree of overlaps can be implemented relatively easily with the GRanges library:

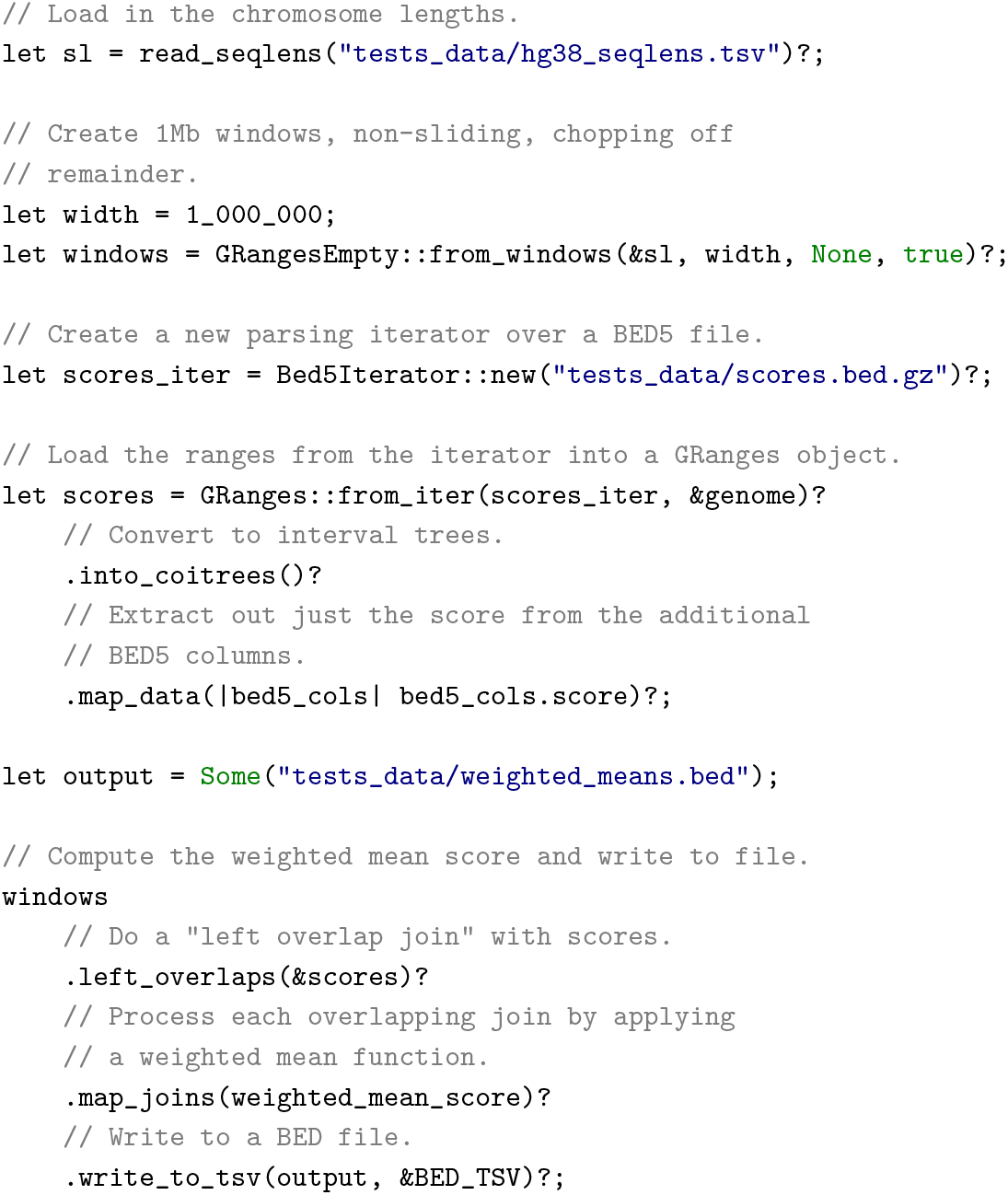

Note that the `weighted mean score()` function that combines the overlapping ranges and their data has been omitted for brevity, but the full example is included in the GRanges library, and can be run with `cargo run --examples weighted mean`. Overall, the GRanges syntax is designed to be expressive yet powerful. This is evident by the fact that the granges command-line tool that accompanies the library implements functionality similar to `bedtools map` in about 70 lines of code. The GRanges library contains parsers for various BED formats, but other bioinformatics formats can be parsed and loaded into a `GRanges<R, T>` object with the Noodles Rust library [11].

### Sequence Types

Often genomic ranges are used to extract the corresponding nucleotide sequences from a genome. The GRanges library generalizes this idea to *any* type of per-basepair data through its `Sequences` trait. For example, a common operation is computing statistical summaries on per-basepair numeric scores, e.g. from CADD [15] or language model effect predictions [2]. The `Sequences` trait provides a common programmatic interface to do such operations, whether computing the variance of pathogenic scores in a window or the GC content of a nucleotide sequence. To avoid unnecessary memory usage, GRanges also supports “lazy” sequence types that access a single chromosome at a time as needed, using the Noodles library [11].

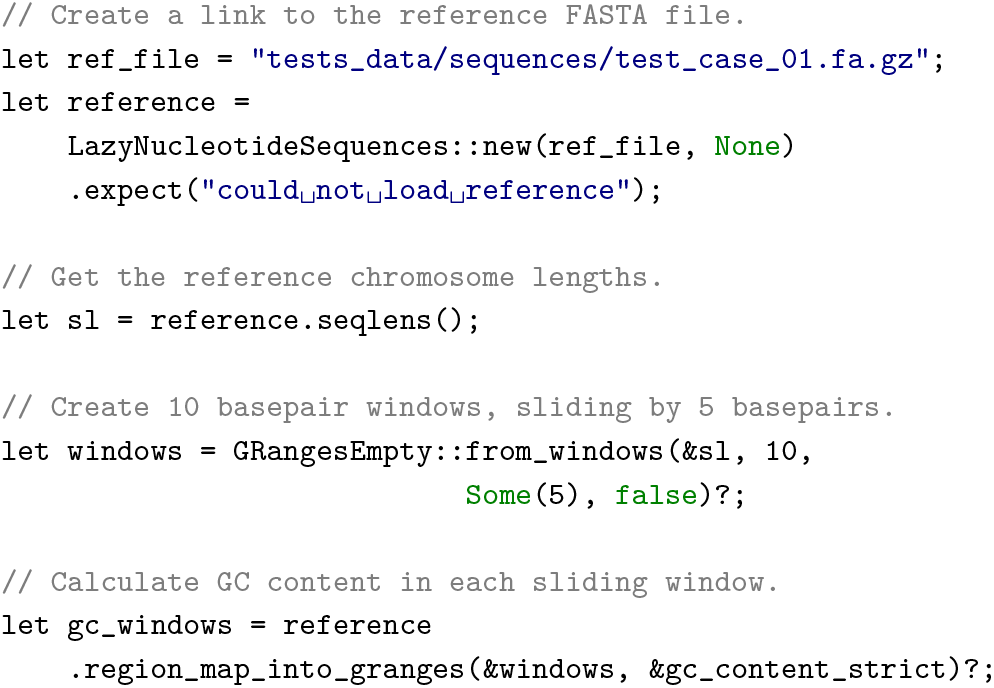

### Performance Comparison

While the primary goal of the GRanges library is to facilitate the development of computational genomics software through its expressive programming interface, because the GRanges library is written in a fast compiled language, the performance of its operations are often faster than comparable operations with bedtools and plyranges (Figure 1). These benchmarks were conducted on a single core of a cluster node with a Intel Xeon Skylake 6130 2.1 GHz processor and 96Gb of memory. The GRanges library also has additional integrated benchmarks against bedtools into the package code to ensure that future developments do not lead to performance regression. While GRanges is faster than bedtools and plyranges, it is about 17% slower than a recently developed Rust-based command-line bedtools alternative, gia [16]. This modest performance cost reflects different specializations: GRanges is not intended to be a high-performance alternative to bedtools like gia or BEDOPs [12], but rather an expressive library for constructing specialized genomic range processing tools.

**Fig. 1.**
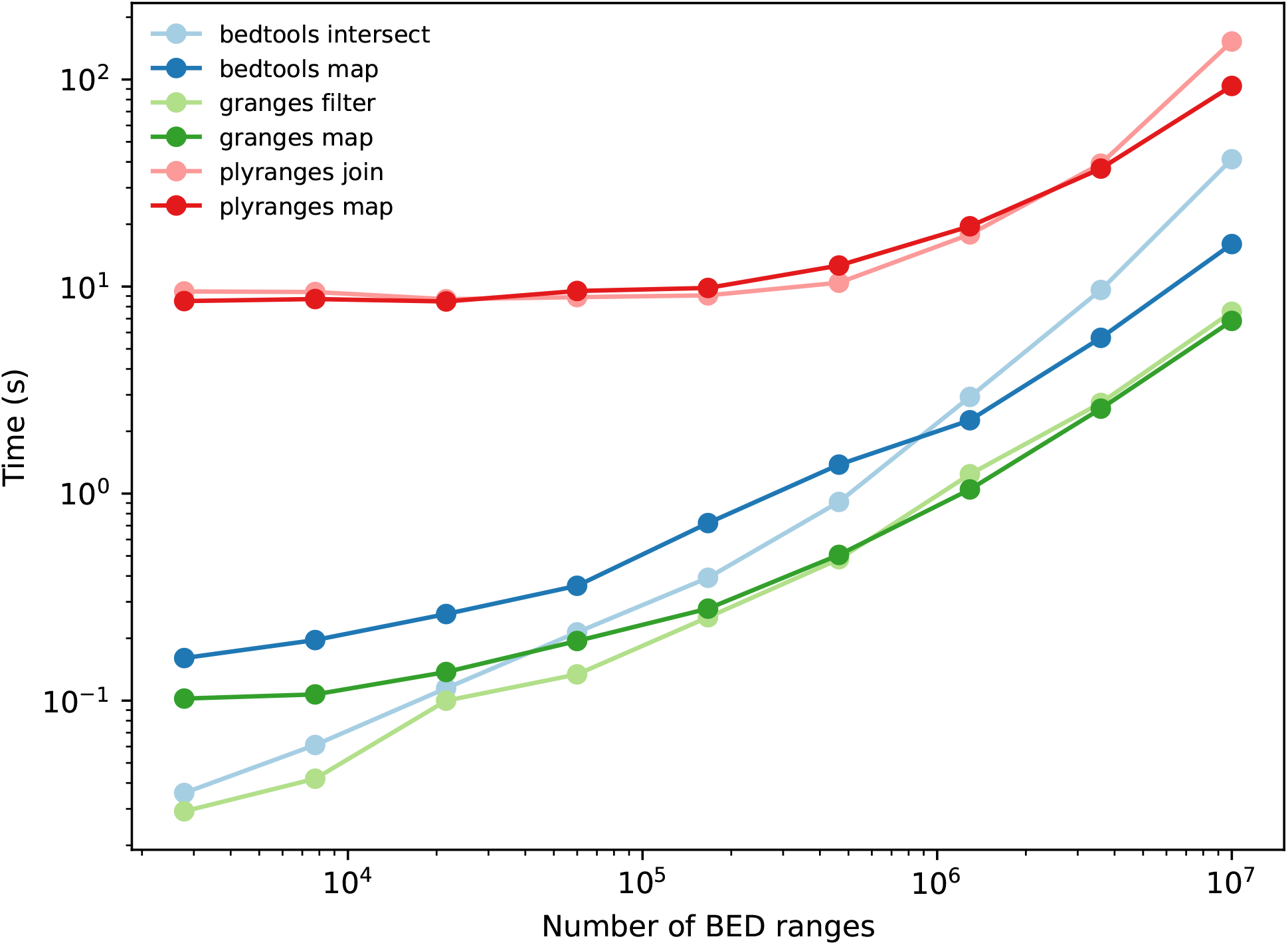
Benchmark of GRange performance compared to bedtools and plyranges for analogous operations.

## Discussion

The GRanges library aims to be a performant and expressive interface to writing software that works with genomic range data. The total library is about 12,200 lines of code, tests, documentation. To safeguard against bugs and numerical inaccuracies, GRanges functions are extensively unit tested and the `granges` command-line tool has extensive integration tests to ensure the output of its commands are identical to the comparable bedtools commands.

Overall, GRanges strikes a balance between flexibility and performance, enabling users to build specialized genomic data processing tools with ease while maintaining high computational efficiency. Furthermore, the simple syntax of the GRanges library will lower the barrier to computational genomics software developers new to the Rust language. The benchmarks illustrate that GRanges outperforms established tools such as bedtools and plyranges in many common genomic range operations. This positions GRanges as a valuable resource for genomic data analysis in the Rust language. Future development on GRanges will focus on expanding the functionality of GRanges to support a wider range of genomic data types and analysis methods, as well as improving performance through multi-threading.

## Data Availability

The GRanges library is available via Rust’s crates system at https://crates.io/crates/granges or on GitHub at https://github.com/vsbuffalo/granges. The library can be added to a Rust project with `cargo add granges`, and the command-line tool can be installed with `cargo install granges`. The documentation for the library is available at https://docs.rs/granges/latest/granges/. GRanges is licensed under the MIT license.

## Author contributions statement

Must include all authors, identified by initials, V.B. conceived of the software and wrote it, and wrote the manuscript.

## Acknowledgments

The author would like to thank Rasmus Nielsen for support during this work, and Rob Patro, Noam Teyssier, and Kevin Thornton for Rust programming advice. The author would also like to thank Daniel C. Jones for developing the superb coitrees library that GRanges uses. This work is supported by NIH grant R01GM138634 awarded to Rasmus Nielsen.

## References

1. M A Bender, E D Demaine, and M Farach-Colton. Cache-oblivious b-trees. In Proceedings 41st Annual Symposium on Foundations of Computer Science, pages 399–409. IEEE, 2000.

2. Gonzalo Benegas, Carlos Albors, Alan J Aw, Chengzhong Ye, and Yun S Song. GPN-MSA: an alignment-based DNA language model for genome-wide variant effect prediction. bioRxiv, April 2024.

3. Wouter De Coster and Rosa Rademakers. NanoPack2: population-scale evaluation of long-read sequencing data. Bioinformatics, 39(5), May 2023.

4. Daniel C Jones. coitrees: A very fast interval tree data structure.

5. Steve Klabnik and Carol Nichols. The Rust Programming Language, 2nd Edition. No Starch Press, February 2023.

6. Johannes Köster. Rust-Bio: a fast and safe bioinformatics library. Bioinformatics, 32(3):444–446, February 2016.

7. Johannes Köster and Sven Rahmann. Snakemake–a scalable bioinformatics workflow engine. Bioinformatics, 28(19):2520–2522, October 2012.

8. Michael Lawrence, Wolfgang Huber, Hervé Pagés, Patrick Aboyoun, Marc Carlson, Robert Gentleman, Martin T Morgan, and Vincent J Carey. Software for computing and annotating genomic ranges. PLoS Comput. Biol., 9(8):e1003118, August 2013.

9. Stuart Lee, Dianne Cook, and Michael Lawrence. plyranges: a grammar of genomic data transformation. Genome Biol., 20(1):4, January 2019.

10. Heng Li. What high-performance language to learn? https://lh3.github.io/2024/03/05/what-high-performance-language-to-learn, March 2024. Accessed: 2024-5-22.

11. Michael Macias. noodles: Bioinformatics I/O libraries. https://crates.io/crates/noodles, May 2024. Accessed: 2024-5-23.

12. Shane Neph, M Scott Kuehn, Alex P Reynolds, Eric Haugen, Robert E Thurman, Audra K Johnson, Eric Rynes, Matthew T Maurano, Jeff Vierstra, Sean Thomas, Richard Sandstrom, Richard Humbert, and John A Stamatoyannopoulos. BEDOPS: high-performance genomic feature operations. Bioinformatics, 28(14):1919–1920, July 2012.

13. Aaron R Quinlan. BEDTools: The Swiss-Army tool for genome feature analysis. Curr. Protoc. Bioinformatics, 47:11.12.1–34, September 2014.

14. Aaron R Quinlan and Ira M Hall. BEDTools: a flexible suite of utilities for comparing genomic features. Bioinformatics, 26(6):841–842, March 2010.

15. Philipp Rentzsch, Daniela Witten, Gregory M Cooper, Jay Shendure, and Martin Kircher. CADD: predicting the deleteriousness of variants throughout the human genome. Nucleic Acids Res., 47(D1):D886–D894, January 2019.

16. Noam Teyssier, Martin Kampmann, and Hani Goodarzi. GIA: A genome interval arithmetic toolkit for high performance interval set operations. September 2023.

17. H Wickham. The split-apply-combine strategy for data analysis. J. Stat. Softw., 2011.

18. H Wickham, M Averick, J Bryan, W Chang, and others. Welcome to the tidyverse. Journal of open source, 2019.

